# Using the Tea Bag Index to determine how two human pharmaceuticals affect litter decomposition by aquatic microorganisms

**DOI:** 10.1101/809426

**Authors:** William Ross Hunter, Ashley Williamson, Judith Maria Sarneel

## Abstract

This study demonstrates that independent additive effects of two human pharmaceuticals, the antibiotic trimethoprim and the artificial estrogen 17a-Ethinylestradiol (EE2), inhibit plant litter decomposition by aquatic microorganisms. The constant release of pharmaceuticals, such as these, has the potential to affect aquatic microbial metabolism and alter biogeochemical cycling of carbon and nutrients. Here we advance the Tea Bag Index (TBI) for decomposition by using it in a series of contaminant exposure experiments testing how interactions between trimethoprim and EE2 affect aquatic microbial activity. The TBI is a citizen science tool used to test microbial activity by measuring the differential degradation of green and rooibos tea as proxies for respectively labile and recalcitrant litter decomposition. Exposure to either trimethoprim or EE2 decreased decomposition of green tea, suggesting additive effects upon microbial activity. Exposure to EE2 alone decreased rooibos tea decomposition. Consequently, trimethoprim and EE2 stabilized labile organic matter against microbial degradation and restricted decomposition. We propose that the method outlined could provide a powerful tool for testing the impacts of multiple interacting pollutants upon microbial activity, at a range of scales, across aquatic systems and over ecologically relevant time scales.

## 1. Introduction

Globally, human pharmaceutical use has increased by 3 % per annum since the year 2000, leading to constant discharge of these compounds into the aquatic environment from both point (Waste Water Treatment Works and Septic Tanks) and diffuse sources (e.g. agricultural run-off) (Gros et al, 2007; Rosi-Marshall & Royer, 2012; Van Boeckel et al, 2014;). Despite being present in the environment at low concentrations, pharmaceuticals have the potential to affect aquatic ecosystem processes because they are designed to be effective at micromolar or nanomolar concentrations (Rosi-Marshall & Royer, 2012; van Broeckel et al., 2014; Álvarez-Muñoz et al. 2015). The specific effects of pharmaceuticals in the environment are likely to be complex as a consequence of the myriad of potential interactions between these compounds, and their potential effects upon aquatic plants, animals and microorganisms (Hernando et al, 2006; Rosi-Marshall & Royer, 2012; Brodin et al, 2014). For example, the presence of antimicrobial compounds has the potential to affect the microbial processes that control aquatic carbon and nitrogen cycling (Rosi-Marshall & Royer, 2012; Brodin et al, 2014; Rosi et al., 2018), whilst other compounds such as artificial estrogens may actually stimulate microbial activity (Ribeiro et al., 2010; Pieper and Rotard, 2011; McClean and Hunter, 2020). As such, we need to understand how the interactions between pharmaceuticals influence microorganism-mediated biogeochemical processes, to safeguard ecosystem services.

Antibiotics and artificial estrogens represent some of the most widely detected pharmaceuticals in the environment (Rosi-Marshall & Royer, 2012; Álvarez-Muñoz et al, 2015; Archer et al, 2017; Gaston et al., 2019). Within aquatic systems, a wide range of human antibiotics such as tetracyclines, sulphonimides and trimethoprim are detectable in the water column, sediments and biota (Metcalfe et al., 2003; Ashton et al., 2004; Thomas and Hilton, 2004, Wiegel et al., 2004; Ebele et al., 2017; Kotke et al., 2019). Trimethoprim is of particular environmental concern, due to its widespread usage in human and veterinary healthcare, where it is mainly used to treat bladder and urinary tract infections, gastro-intestinal problems, ear infections and as prohylaxis against opportunistic infections (World Health Organisation, 2019). Approximately 50 – 60 % of any administered trimethoprim dose is excreted in a subject’s urine, combined with its widespread use this provides a definite route into the environment via municipal wastewater and leaching of agricultural effluents. As such, alongside its categorization as an essential medicine (World Health Organisation, 2019), trimethoprim is recognized as a potential environmental hazard (Straub, 2013; Tell et al., 2019).

The presence of antibiotics and their residues, both in the water column and aquatic sediments may, directly affect the structure (Wieser et al 2016; Yuan et al 2017) and functioning of aquatic microbial communities. In inland waters, leaf litter and other terrigenous phytodetritus represent an important energy and nutrient source within the food-web (e.g. Vannote et al, 1980; Raymond & Bauer, 2001; Raymond et al., 2016). Microbial decomposition is an essential process, which supports the transfer of litter-derived energy and nutrients to organisms at higher trophic levels. Consequently, antimicrobials are likely to directly affect the processes that heavily depend on microbial communities, and alter ecosystem-scale carbon and nitrogen cycling will be affected (Roose-Amsleg and Laverman, 2016; Robson et al., 2020). Furthermore, whilst bacterial populations rapidly evolve antibiotic resistance, this comes at a physiological cost. Antibiotic resistant bacteria tend to grow more slowly and use nutrients less efficiently than their non-resistant conspecifics (Anderson and Levin, 1999, Jansen et al 2013). As such, longer-term exposure to antimicrobial contaminations is likely to have implications for the efficiency of microbially-mediated processes in aquatic ecosystems (e.g. Rosi et al. 2018).

By contrast, artificial estrogens, such as EE2, may provide a novel carbon source for aquatic microorganisms, supporting increased microbial metabolism (Ribeiro et al., 2010; Pieper and Rotard, 2011), their adsorption onto microbial biofilms can provide protection against antimicrobial agents (Writer et al., 2012; Zhang et al., 2014).This was recently demonstrated by McClean and Hunter (2020), who showed that exposure of streambed biofilms to EE2 counteracted the inhibitory effects of ibuprofen upon microbial respiration. Consequently, the interactions between antibiotics and artificial estrogens upon aquatic microbial processes are likely to have wider environmental relevance.

Contaminant exposure substrata (CES) experiment provides a powerful tool for testing the sensitivity of aquatic microbial communities to pharmaceuticals and other pollutants (Tank et al, 2006; Costello et al, 2016). A well-refined method is provided by Costello et al (2016) to test how pharmaceuticals affect microbial biofilm growth and community structure and ecophsysiological responses of the biofilm, such as community respiration (Rosi-Marshall et al, 2013; Rosi et al., 2018; Gallagher and Reisinger, 2020; McClean & Hunter, 2020 Robson et al., 2020). These methods, however, provide insufficient information on heterotrophic processes such as microbial litter decomposition, which occurs over times-scales measured in weeks or months (Vannote et al, 1980; Raymond & Bauer, 2001). The Tea Bag Index (TBI) provides a powerful low-cost tool for investigating microbial activity in soils and aquatic systems, based upon the traditional use of leaf-litter bags in ecology (Keuskemp et al., 2013; Whigham et al., 2017; Mueller et al., 2018; Seelen et al., 2019. The key strength of TBI is its ability to provide a standardized method of quantifying microbial activity by comparing the relative degradation of green tea (easy to decompose, or labile material) and rooibos tea (resistant to decomposition, recalcitrant, lignified material) organic matter source. The use of TBI within contaminant exposure substrata experiments will allow quantification of the impacts of pollutant exposure on microbially-mediated litter decomposition.

Within this study we investigate how interactions between the antibiotic trimethoprim and the artificial estrogen EE2 affect aquatic litter decomposition, using modified contaminant exposure experiments. We integrate the TBI (Keuskemp et al., 2013) into the existing CES methodology developed by Tank et al (2006) and Costello et al (2016). This will provide a simple, low-cost tool for assessing how pharmaceuticals (and other contaminants) may affect microbial litter decomposition. Using the TBI-CES method we test how exposure of the microbial community to both trimethoprim and EE2 affects the decomposition of both labile and recalcitrant litter in a mixed-use catchment stream. We hypothesize that trimethoprim will inhibit decomposition while EE2 may enhance it. We therefore expect that EE2 may mediate against any inhibitory effects of trimethoprim upon the microbial activity that drives litter decomposition.

## 2. Materials and Methods

### 2.1. Site Description

Experiments were carried out between 22^nd^ March and 13^th^ June 2019 in the Ballysally Blagh (Latitude: 55°08’48.9”N Longitude: 6°40’08.0”W), a ground-water fed second-order stream draining a mixed agricultural (consisting of 21.9 % arable; 55.9 % grassland; 13.7 % heathland; 1.9 % woodland) and urban (7.3 %) catchment of 14.2 km^2^. Mean volumetric water flow in the Ballysally Blagh is 0.21 (± 0.27) m^3^ s^-1^, measured at a V-shaped weir. Water temperatures at the study site (50 cm water depth) were recorded at 1-hour intervals throughout the experiment using a HOBO MX2204 Bluetooth temperature logger and ranged between 5.51 °C and 15.57 °C, with a mean temperature over the study period of 10.17 (± 2.08) °C.

### 2.2. Contaminant Exposure Substrata Experiments

Contaminant exposure substrata (CES) experiments were conducted using forty 120 ml screw cap sample pots, with a 35 mm hole drilled into the lid (following Costello et al, 2016). Each pot was filled with 100 ml of 2 % agar, with ten pots spiked with 688 μmol.l^-1^ Trimethoprim (Sigma-Aldich, product no: 92131), ten with 688 μmol.l^-1^ of EE2 (Sigma-Aldrich, Product No. E4876) and ten with 688 μmol.l^-1^ trimethoprim and 688 μmol.l^-1^ EE2. The remaining ten sample pots received no pharmaceutical treatment (control). Trimethoprim and EE2 stock solutions for were made up in 70 % ethanol, with 1 ml aliquots used to spike each contaminant exposure experiment, whilst the agar in the control treatments was spiked with 1 ml doses of 70 % ethanol. All pots were weighed to determine mass of agar within. One pre-weighed unused Lipton® Green Tea Sencha pyramidal tea bag with non-woven fabric (Product Number: EAN 87 22700 05552 5) and one pre-weighed unused Lipton® Rooibos tea bag with non-woven fabric (Product Number: EAN 87 22700 18843 8) were the placed into each pot screw cap secured in place. The contaminant exposure experiments were then secured to four L-shaped metal bars (Length = 1000 mm; Width = 35 mm; Height = 35 mm) and deployed onto the streambed at 30 – 40 cm depth (Figure 1).

**Figure 1.**
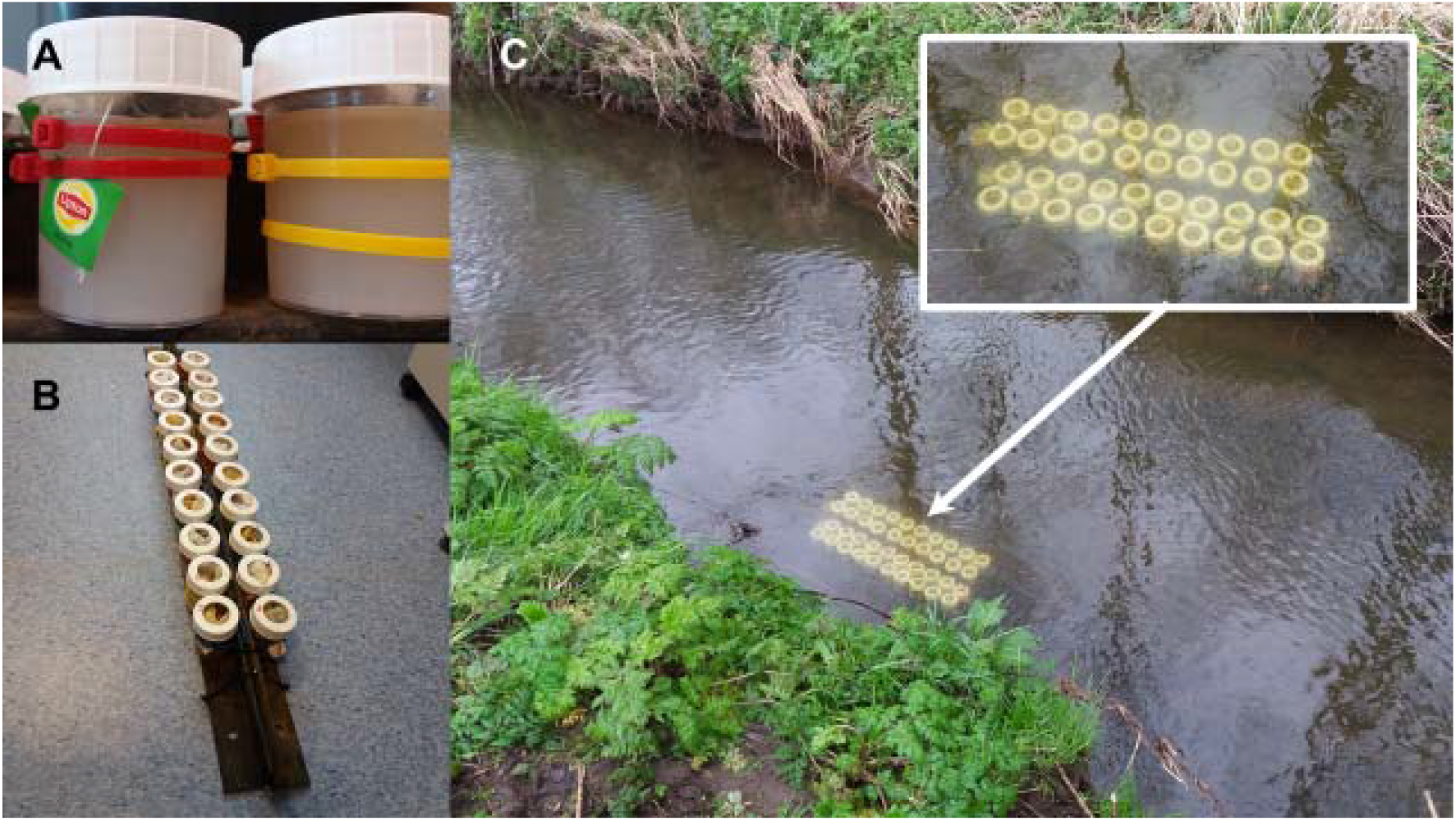
Images of (A) the TBI contaminant exposure experiments, (B) how they are secured prior to the L-shaped metal bars and (C) their deployment at the streambed (right panel)

CES experiments were left in place for 83 days. Upon recovery, the teabags were removed from the experiment and the pots were then weighed to determine the agar mass loss (following Costello et al, 2016). Teabags were dried to constant weight over 72 hours at 60 °C, excess silt and detritus adhering to each teabag was gently removed, and their dry-weight determined (following Keuskamp et al, 2013). The tea was then removed from each teabag, re-weighed and combusted at 550 °C for four hours to determine ash free dry mass.

### 2.3. Data Treatment

We calculated percentage mass loss of green tea and rooibos tea in each contaminant exposure experiment as the difference between the dry weight of the teabag at the start and end of the incubation. We corrected for mass loss during the drying process based on the mean mass loss for five air-dry green teabags and five air-dry rooibos teabags, dried at 60 °C over 72 hours. Following Keuskamp et al. (2013) we then determined the organic matter stabilization factor (*S*_*TBI*_) as:

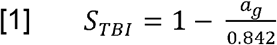

Where *a*_*g*_ is the decomposable fraction of the green tea (ash free dry weight) and 0.842 is the hydrolysable fraction of the green tea (from Keuskamp et al, 2013). Using S we then estimated the decomposable fraction of the rooibos tea (*a*_*r*_) as:

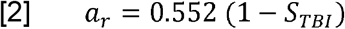

Where *S*_*TBI*_ is the stabilization factor and 0.552 is the hydrolysable fraction of the rooibos tea (from Keuskamp et al, 2013). The organic matter Initial decomposition rate of the labile material (*k*_*TBI*_) can then be calculated as:

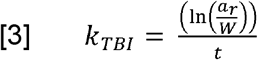

Where *W* is the mass loss of the Rooibos teabag in each experiment (in grams) and *t* is the duration of the experimental incubation (in days).

### 2.4. Data Analysis

Data analysis and visualization was carried out in R using the *base* and *ggplot* packages (R Development Core Team, 2009; Wickham, 2016). We tested for independent and combined effects of trimethoprim and EE2 upon the percentage mass loss of the Green tea and Rooibos teabags, the stabilization factor (*S*_*TBI*_) and Initial decomposition rate (*k*_*TBI*_) of the labile organic material using two-way analysis of variance (ANOVA). Post-hoc testing of significant interactions were conducted using Tukey’s test for Honest Significant Difference. All data were visually explored to ensure they conformed to the assumptions of normality and homoscedacity (following Zuur et al, 2010), Initial decomposition rate (*k*_*TBI*_) data were log_10_ transformed to ensure that the residuals from the ANOVA model conformed to a normal distribution. See supplementary table 1 for a summary of the ANOVA test results.

### 3. Results and Discussion

Over the course of the experiment, dissolution of the agar was assumed to deliver constant daily doses of approximately 275 nmol. d^-1^ of trimethoprim and/or EE2 in both the single and combined pharmaceutical treatments. As such, each teabag would have been exposed to a pharmaceutical dose that was broadly comparable with the concentration of these compounds in inland waters (e.g. Álvarez-Muñoz et al, 2015; Archer et al, 2017; Tousova et al., 2017). Rooibos Tea mass loss decreased upon EE2 exposure, with no additional/significant effects of trimethoprim (Figure 2 A, Table 1a). By contrast for green tea, we observed additive and inhibiting effects of both trimethoprim and EE2 (Figure 2 B, Table 1b). Based on these data we can demonstrate that EE2 and trimethoprim had significant independent effects upon both the TBI stabilization factor (Figure 3 A, Table 1c) and initial decomposition rate (Figure 3 B, Table 1 d).

**Figure 2.**
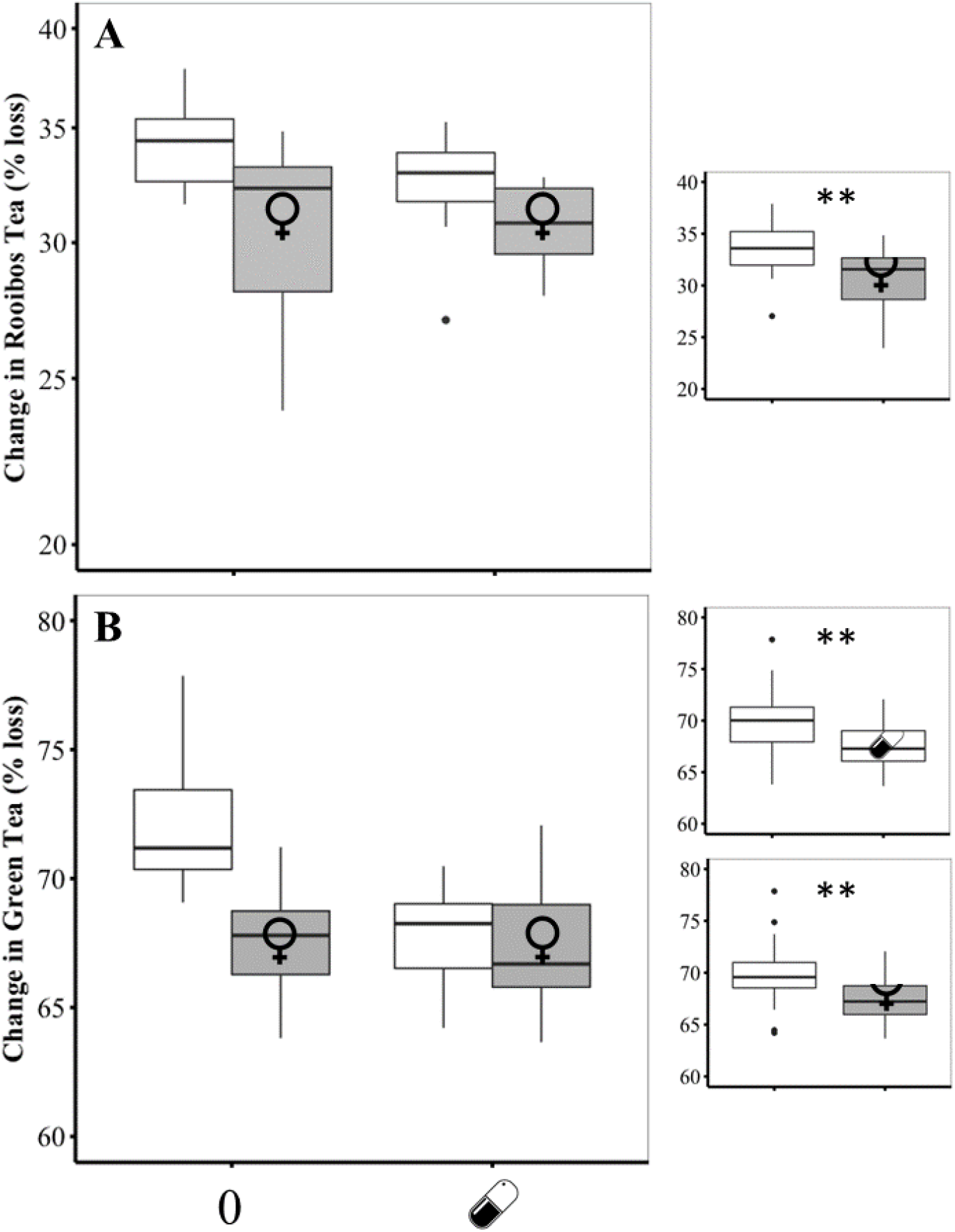
Effects of TMP 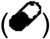 and EE2 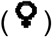 on the degradation (% mass loss) of A) Rooibos and B) Green tea, as labile and recalcitrant organic matter sources. Inserts show pooled data where significant independent effects of either Trimethoprim or EE2 where detected. Significance levels: *** *p* < 0.001; ** *p* < 0.01; * *p* < 0.05.

**Figure 3.**
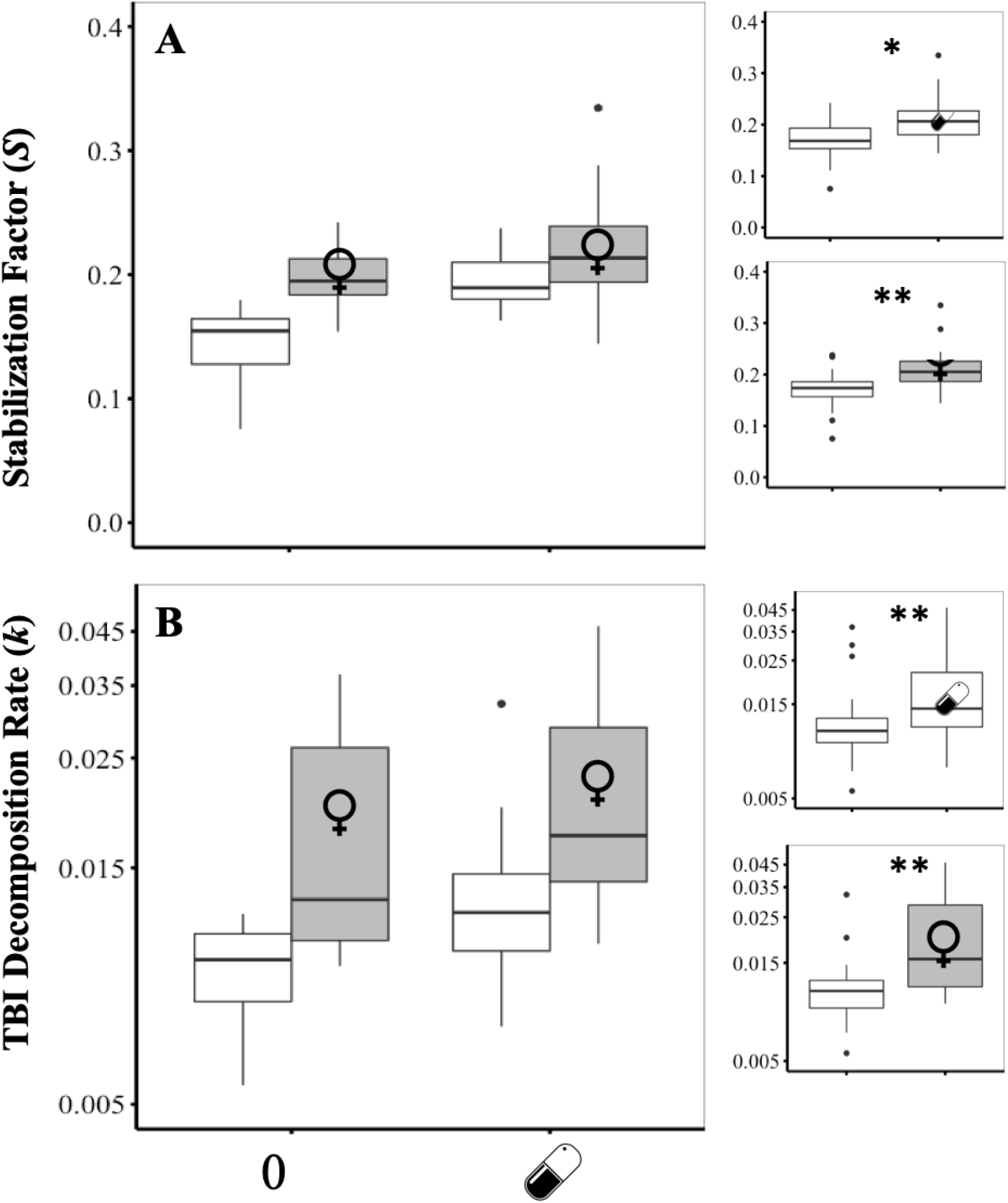
Effects of TMP 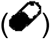 and EE2 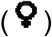 on the A) TBI stabilization factor and B) initial decomposition rate, calculated following Keuskamp et al (2013). Inserts show pooled data where significant independent effects of either Trimethoprim or EE2 where detected. Significance levels: *** *p* < 0.001; ** *p* < 0.01; * *p* < 0.05.

**Table 1.**
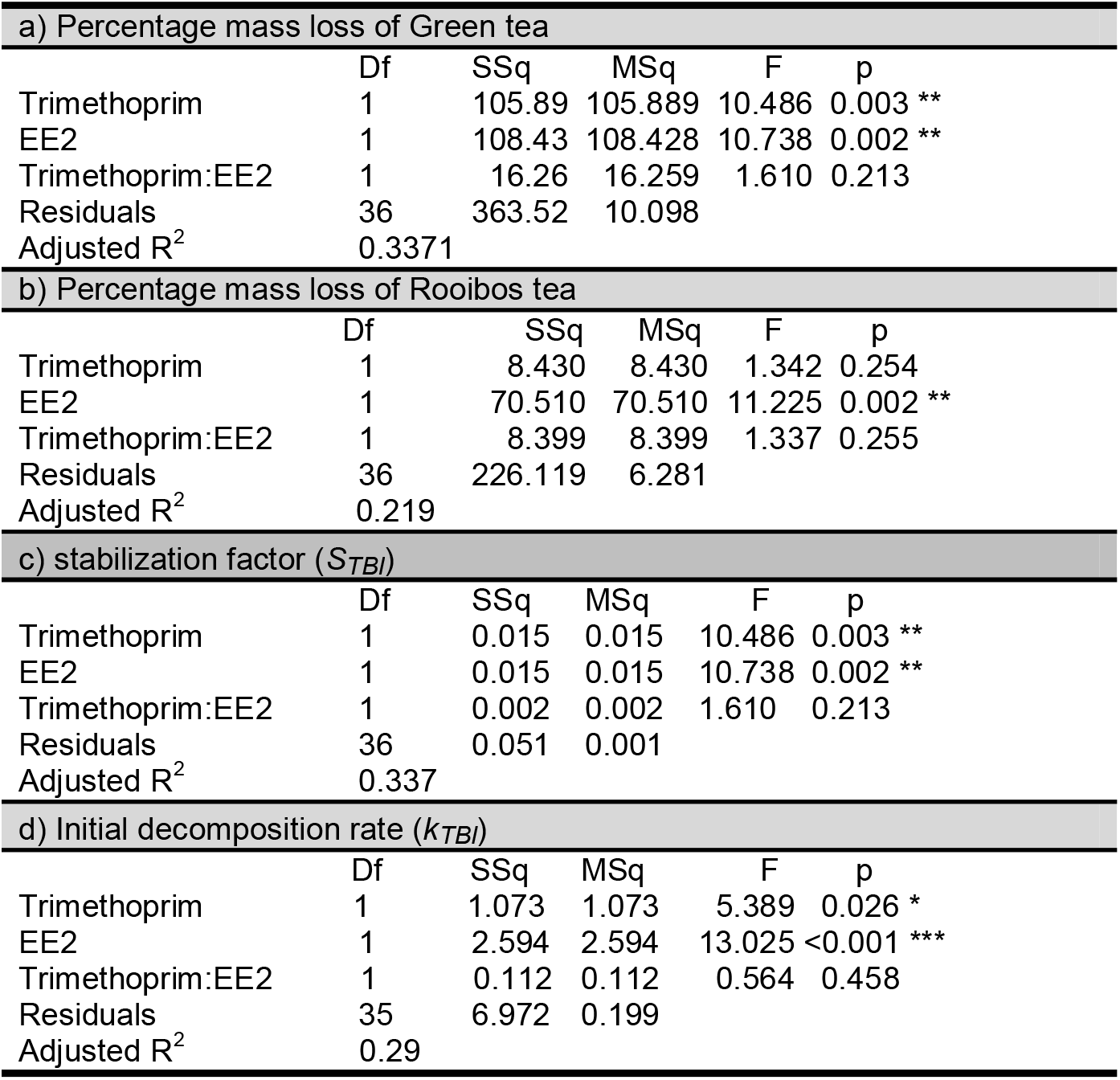
Two-way analysis of variance testing for significant effects of trimethoprim and EE2 upon the percentage mass loss of the Green tea and Rooibos teabags, the organic matter stabilization factor (*S*_*TBI*_) and initial decomposition rate (*k*_*TBI*_).

| a) Percentage mass loss of Green tea |  |  |  |  |  |
| --- | --- | --- | --- | --- | --- |
|  | Df | SSq | MSq | F | p |
| Trimethoprim | 1 | 105.89 | 105.889 | 10.486 | 0.003 ** |
| EE2 | 1 | 108.43 | 108.428 | 10.738 | 0.002 ** |
| Trimethoprim:EE2 | 1 | 16.26 | 16.259 | 1.610 | 0.213 |
| Residuals | 36 | 363.52 | 10.098 |  |  |
| Adjusted R<sup>2</sup> | 0.3371 |  |  |  |  |
| b) Percentage mass loss of Rooibos tea |  |  |  |  |  |
|  | Df | SSq | MSq | F | p |
| Trimethoprim | 1 | 8.430 | 8.430 | 1.342 | 0.254 |
| EE2 | 1 | 70.510 | 70.510 | 11.225 | 0.002 ** |
| Trimethoprim:EE2 | 1 | 8.399 | 8.399 | 1.337 | 0.255 |
| Residuals | 36 | 226.119 | 6.281 |  |  |
| Adjusted R<sup>2</sup> | 0.219 |  |  |  |  |
| c) stabilization factor ($S_{TBI}$) | | | | | |
|  | Df | SSq | MSq | F | p |
| Trimethoprim | 1 | 0.015 | 0.015 | 10.486 | 0.003 ** |
| EE2 | 1 | 0.015 | 0.015 | 10.738 | 0.002 ** |
| Trimethoprim:EE2 | 1 | 0.002 | 0.002 | 1.610 | 0.213 |
| Residuals | 36 | 0.051 | 0.001 |  |  |
| Adjusted R<sup>2</sup> | 0.337 |  |  |  |  |
| d) Initial decomposition rate ($k_{TBI}$) | | | | | |
|  | Df | SSq | MSq | F | p |
| Trimethoprim | 1 | 1.073 | 1.073 | 5.389 | 0.026 * |
| EE2 | 1 | 2.594 | 2.594 | 13.025 | <0.001 *** |
| Trimethoprim:EE2 | 1 | 0.112 | 0.112 | 0.564 | 0.458 |
| Residuals | 35 | 6.972 | 0.199 |  |  |
| Adjusted R<sup>2</sup> | 0.29 |  |  |  |  |

Trimethoprim decreased mass loss of green tea but not of rooibos, while EE2 decreased both green tea and rooibos mass loss. Together, these pharmaceuticals had additive negative effects on green tea mass loss but not on rooibos. No significant interaction was observed for either green tea or rooibos. Further, EE2 and trimethoprim had significant independent effects upon both the litter stabilization factor (Figure 3 A, Table 1c) and initial decomposition rate (Figure 3 B, Table 1 d)

Our results demonstrate that exposure to low doses of both an antibiotic (trimethoprim) and an endocrine disruptor (EE2) inhibit microbial litter decomposition. Specifically, trimethoprim appears to affect decomposition of lignified material less compared to more easy to degrade material, whilst EE2 had a broader effect across litter types. In counter to our hypothesis, EE2 did not reduce the negative impact of trimethoprim upon aquatic litter decomposition. Both trimethoprim and EE2 had significant additive positive effects upon the organic matter stabilization factor (*S*_*TBI*_) and its initial decomposition rate (*k*_*TBI*_). Our study, thus, indicates that pollution from both pharmaceuticals may increase the residence time for organic matter in aquatic systems. Inland waters (rivers, lakes and streams) typically receive large inputs of terrestrial organic matter, which is then partially metabolized, temporarily buried within the sediment or transported to the ocean (Battin et al, 2009). Our study adds further evidence to that pharmaceutical contaminants directly affect the rates and pathways for aquatic carbon and nutrient cycling (e.g. Jobling et al. 2003; Hernando et al. 2006; Gros et al. 2007; Rosi-Marshall and Royer 2012; Rosi-Marshall et al. 2013; Álvarez-Muñoz et al. 2015; Ruhí et al. 2016; Archer et al. 2017; Rosi et al. 2018; Gallagher and Reisinger 2020; McClean and Hunter, 2020).

Given the widespread use of trimethoprim as an antibiotic medication and its high solubility in water [ranging from 500 mg l^-1^ at pH 8 to 15,500 mg l^-1^ at pH 5.5 (Dahlan et al. 1987)], its effect upon microbial activity within the teabag experiment was expected. As a highly soluble contaminant trimethoprim is likely to leach rapidly from the agar within CES experiments, and thus affect microbial activity during the early phases of the experiment. Thus the effects of trimethoprim upon labile litter decomposition are likely to reflect the relatively high solubility of this contaminant. By contrast, EE2 exhibits relatively low solubility in water, ranging between 2.92 and 4.8 mg l-1 (Lai et al., 2000; Ying et al. 2002; Guo and Hu, 2014). This property of EE2, combined with its tendency to adsorb onto organic substances (Lai et al., 2000; Ying et al., 2002; Writer et al., 2012; Zhang et al., 2014), provides a mechanism through which the EE2 released by the agar would affect the degradation of more recalcitrant and lignified litter. The additive effects of trimethoprim and EE2 upon litter decomposition within our experiment were likely to be driven by differences in the solubility of each compound and their potential to adsorb onto the tea. This highlights how differences in the physico-chemical properties of pharmaceutical mediate their potential impacts in aquatic biogeochemical processes.

CES experiments are typically used to determine the effects of a selected contaminant on biofilm forming bacteria and algae (Tank et al., 2006; Costello et al., 2016). Integrating the TBI (Keuskemp et al. 2013) into CES experiments provides a useful low-cost tool to quantify how exposure to pollutants such as pharmaceuticals affects instream decomposition of organic matter. The inclusion of both a labile (green tea) and more recalcitrant (rooibos tea) organic matter source within the experiment allows inferences to made upon the effects of a contaminant upon the phenology of the litter decomposition process. Refinement of the method is, however, necessary and should include investigation of agar dissolution and teabag degradation rates, and how these may vary under differing climatic and hydrological regimes (following Costello et al., 2016).

Whilst our study was restricted to one stream, the method could easily be replicated at multiple sites, within more complex experimental designs. We believe this method will allow the effects of multiple pollutants upon organic matter degradation and residence times to be made across a range of spatial and temporal scales in both freshwater and marine systems. As our method is characterized by its low-cost, simplicity and use of readily available consumer products, it could potentially be used to achieve high levels replications within experimental designs testing the interactions between a suite of multiple pollutants and their effects upon aquatic biogeochemical processes. Furthermore, we propose that the TBI-CES method could provide a valuable pedagogical tool for research-led environmental science teaching and in citizen science based initiatives to investigate the impacts of pollutants in aquatic systems.

## Acknowledgements

This work was funded through start-up funds provided to WRH by the University of Ulster School of Geography and Environmental Science. JMS acknowledges the Swedish Research Council (Vetenskapsrådet) for funding.

## Author Information

Experiments were designed by WRH following discussions with JS, and were conducted by WRH and AW. All authors contributed to the writing of the manuscript.

## Conflict of Interest

The authors declare that we have no conflicts of interest pertaining to this study.

## Data Availability

All data are freely available Creative Commons (CC-BY) license through the Zenodo data repository (DOI: 10.5281/zenodo.3528009).

## Animal Research (Ethics)

No animals were used in this research.

## Consent to Participate (Ethics)

No human subjects were used in this research.

## Consent to Publish (Ethics)

This is not applicable to this study.

## Plant Reproducibility

This is not applicable to this study.

## Clinical Trials Registration

This is not applicable to this study.

## Notes

https://zenodo.org/record/3528009#.X6qbj1pxfcs

### Competing Interest Statement

The authors have declared no competing interest.

### Summary of Updates

Updated manuscript structure to include a detailed materials and methods section and some additional discussion / interpretation. Changes to the title to better reflect the focus of the paper. More detail added to the introduction.

